# Cyclin B3 is specifically required for metaphase to anaphase transition in mouse oocyte meiosis I

**DOI:** 10.1101/390351

**Authors:** Yufei Li, Leyun Wang, Linlin Zhang, Zhengquan He, Guihai Feng, Hao Sun, Jiaqiang Wang, Zhikun Li, Chao Liu, Jiabao Han, Junjie Mao, Xuewei Yuan, Liyuan Jiang, Ying Zhang, Qi Zhou, Wei Li

## Abstract

Meiosis, a cell division to generate gametes for sexual reproduction in eukaryotes, executes a single round of DNA replication and two successive rounds of chromosome segregation [1]. The extraordinary reliability of the meiotic cycle requires the activities of cyclin-dependent kinases (Cdks) associated with specific cyclins [2-4]. Cyclins are the regulatory subunits of protein kinases, which are the main regulators of maturation promoting factor or mitosis promoting factor (MPF) [5, 6] and anaphase-promoting complex/cyclosome (APC/C) [7, 8] in eukaryotic cell division. But how cyclins collaborate to control meiosis is still largely unknown. Cyclin B3 (Ccnb3) shares homology with A- and B-type cyclins [9], and is conserved during higher eukaryote evolution [10-17]. Previous studies have shown that *Ccnb3*-deleted females are sterile with oocytes unable to complete meiosis I in *Drosophila* [18], implying that Ccnb3 may have a special role in meiosis. To clarify the function of Ccnb3 in meiosis in mammalian species, we generated *Ccnb3* mutant mice by CRISPR/Cas9, and found that *Ccnb3* mutation caused female infertility with the failure of metaphase-anaphase transition in meiosis I. Ccnb3 was necessary for APC/C activation to initiate anaphase I, but not required for oocytes maturation, meiosis II progression, or early embryonic development. Our study reveals the differential cell cycle regulation between meiosis I and meiosis II, as well as meiosis between males and females, which shed light on the cell cycle control of meiosis.

**Highlights:**

- Identification of a female meiosis-specific cyclin in mouse
- Cyclin B3 is required for metaphase-anaphase transition in oocyte meiosis I
- Cyclin B3 is not essential for oocyte maturation and sister chromosome segregation
- Cyclin B3 is necessary for APC/C activation and MPF kinase activity through Cdk1

## Results

### *Ccnb3* mutation leads to female infertility

We firstly detected the expression pattern of Ccnb3 by quantitative PCR (Q-PCR) and found that its mRNA had similar expression pattern with cyclin B1 (Ccnb1) during oocyte *in vitro* maturation, which implied that Ccnb3 may play an important role in meiosis cell cycle regulation (Figure 1A). To study the role of Ccnb3, we generated the *Ccnb3* mutant mice (referred to as *Ccnb3*^Δ/Y^ and *Ccnb3*^Δ/Δ^ for the male and female mutants respectively) via CRISPR-mediated deletion of 29 base pair (bp) in exon 3 of *Ccnb3* gene located on the X chromosome (Figure S1A). The genotype of *Ccnb3* mutant mice were verified by PCR (Figure S1B). By natural mating, we found that the *Ccnb3*^Δ/Δ^ female mice were infertile, while the *Ccnb3*^Δ/Y^ male mice showed normal fertility (Figure 1B). To find the cause of female infertility, we examined the ovary development and folliculogenesis in *Ccnb3*^Δ/Δ^ female mice. The hematoxylin-eosin (H&E) staining results showed that the *Ccnb3*^Δ/Δ^ ovary development was normal (Figure 1C), and the number of superovulated oocytes of *Ccnb3*^Δ/Δ^ mice were similar with the wild type female mice (referred to as *Ccnb3*^WT/WT^) (Figure S1C). To investigate whether the infertility was caused by embryonic lethality, we collected embryos from *Ccnb3*^Δ/Δ^ female mice with vaginal plugs after mating with *Ccnb3*^WT/Y^ male mice. All the collected fetuses were degenerated before embryonic day 7.5 (E7.5) (Figure 1D). These results demonstrated that *Ccnb3* mutation leads to female infertility, while the defects were caused by embryonic lethality rather than the abnormal follicular development.

**Figure 1.**
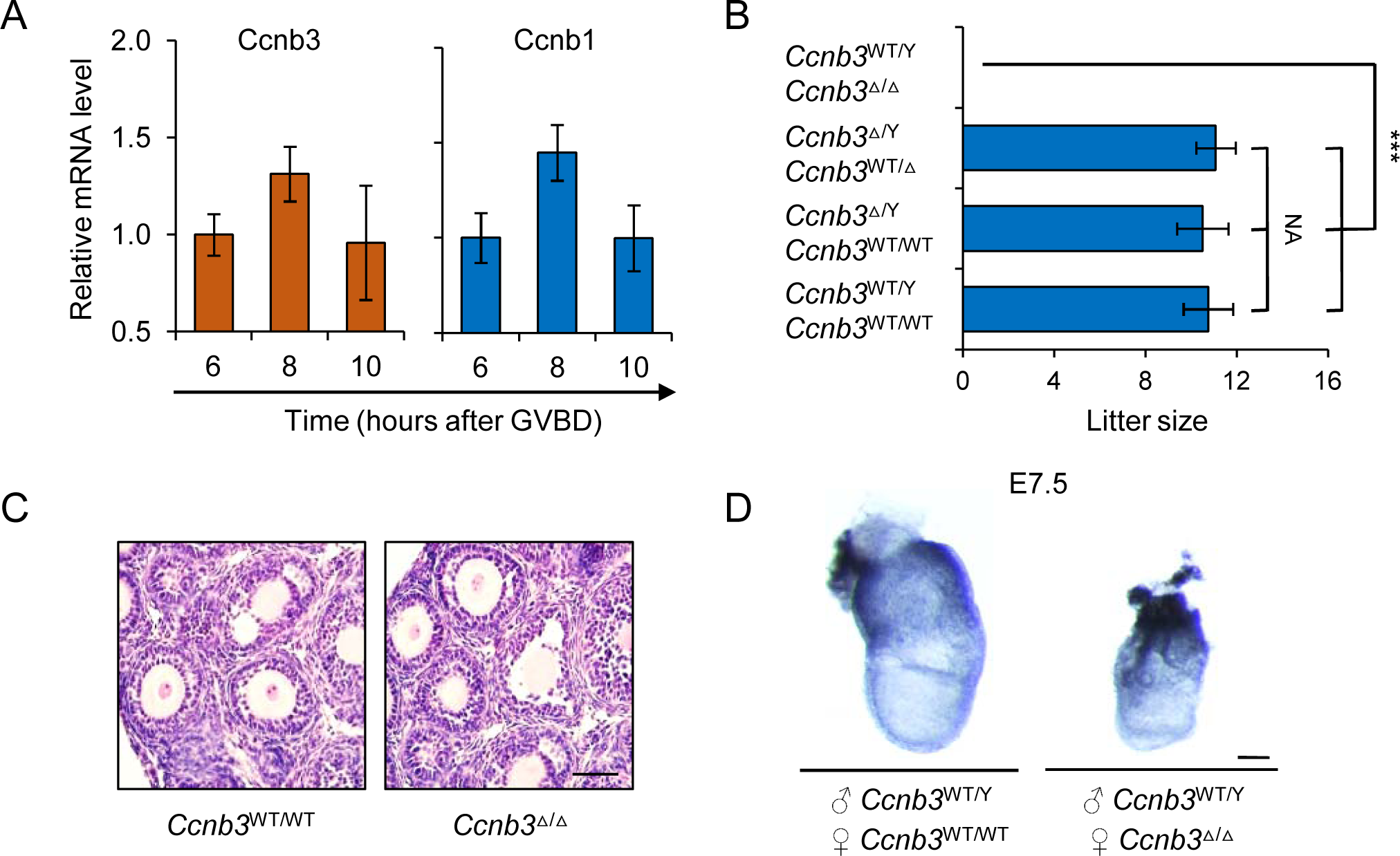
*Ccnb3* mutation led to female infertility in mice. (A) The mRNA expression pattern of Ccnb1 and Ccnb3 in mouse oocytes during *in vitro* maturation (n = 40 in each group). (B) Litter size counts showing that the *Ccnb3*^Δ/Δ^ mice mated with wild type male mice failed to produce full-term offspring. Unpaired two-tailed Student’s *t*-test. Error bars represent mean ± SD. ***p < 0.001. (C) H&E staining of *Ccnb3*^Δ/Δ^ ovary. The morphology of ovary and the number of follicles in *Ccnb3*^Δ/Δ^ mice were similar with that in the wild type mice. Scale bar, 40 μm. (D) The embryos produced by mating *Ccnb3*^Δ/Δ^ mice with wild type male mice was died before embryonic day 7.5 (E7.5).

### *Ccnb3* mutation causes oocyte meiotic arrest at metaphase I (MetI)

Although the number of superovulated oocytes from *Ccnb3*^Δ/Δ^ female mice was identical to that from *Ccnb3*^WT/WT^ mice, the first polar body (PB) were not observed in the *Ccnb3*^Δ/Δ^ oocytes (Figure 2A). We suspected that Ccnb3 loss-of-function may lead to defects during meiosis progression. To confirm this hypothesis, we analyzed the maturation process of the *Ccnb3*^Δ/Δ^ oocytes using *in vitro* maturation (IVM) (Figure 2B) and living cell tracking assays. We found that the fully grown germinal vesicle (GV)-stage *Ccnb3*^Δ/Δ^ oocytes could resume meiosis, break down germinal vesicles (GVBD), and form metaphase I (MetI) spindles, suggesting that *Ccnb3* mutation did not affect the GVBD efficiency of *Ccnb3*^Δ/Δ^ oocytes (81.1% *vs* 88.5%) and further development into MetI stage (Figure 2C and 2D, Figure S1D and S1E). However, after further culturing the oocytes to time point corresponding to metaphase II (MetII) stage, the *Ccnb3*^Δ/Δ^ oocytes failed to extrude the first polar body (PB) and still maintained bivalent homologous chromosomes, when the *Ccnb3*^WT/WT^ oocytes had entered to MetII stage with first PB and univalent sister chromatids (Figure 2D and 2E). These results showed that *Ccnb3* mutant oocytes failed to segregate homologous chromosomes and arrested at MetI stage. The Rec8 protein, a specific meiotic cohesion, was still present on the chromosome arm in the *Ccnb3*^Δ/Δ^ oocytes arrested at MetI, indicating that the homologous chromosomes were unable to disjoin in *Ccnb3*^Δ/Δ^ oocytes (Figure 2F).

**Figure 2.**
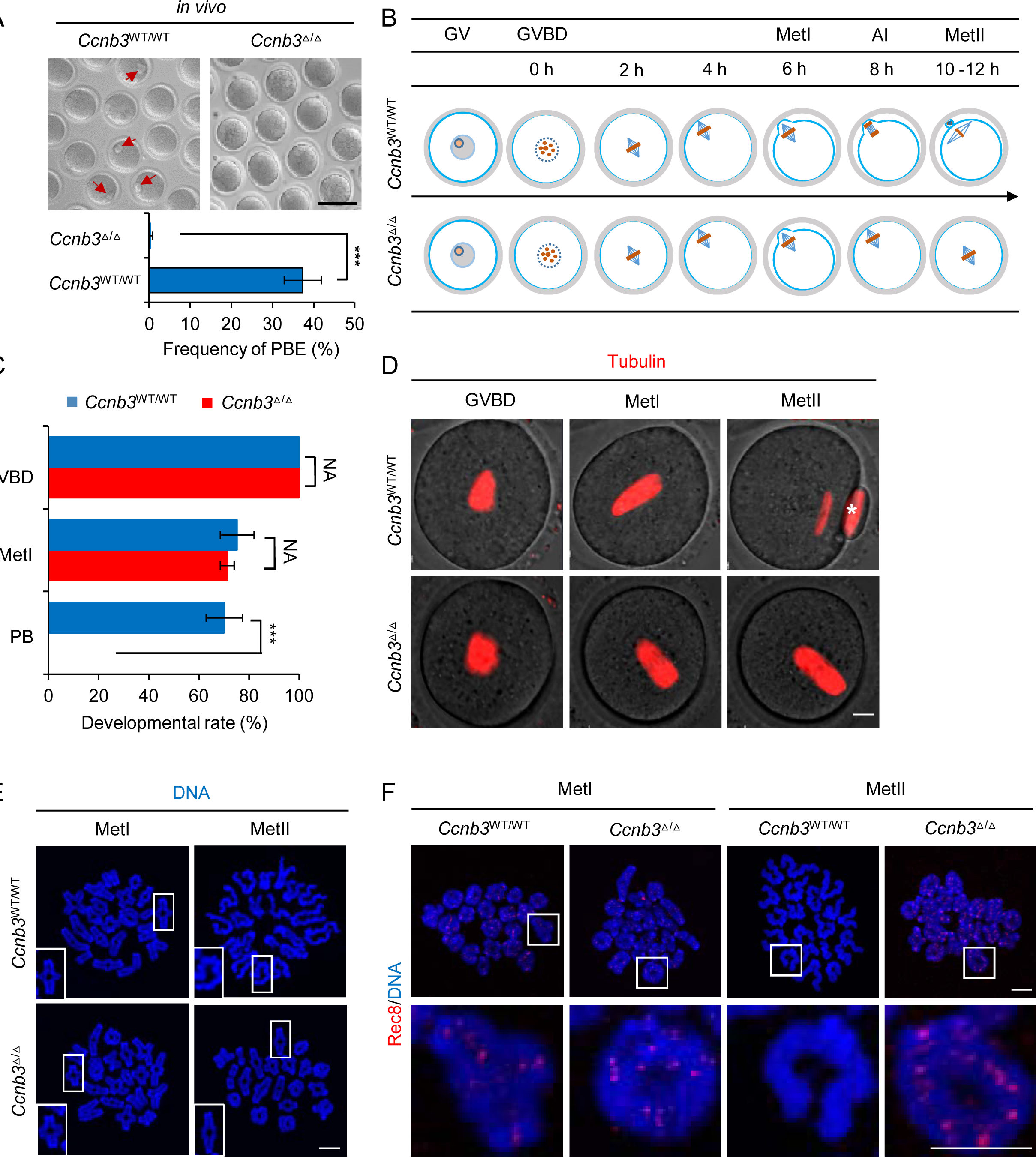
*Ccnb3* mutation caused mouse oocyte meiotic arrest at metaphase I. (A) Oocytes with *Ccnb3* mutation failed to extrude first polar body (PB) by *in vivo* superovulation. Red arrow heading to the first PB. Unpaired two-tailed Student’s *t*-test. Error bars represent mean ± SD. ***p < 0.001. (B) Schematic outline of the *in vitro* maturation process of mouse oocytes. (C) Oocytes with *Ccnb3* mutation failed to extrude first PB by *in vitro* maturation assay. Oocytes were incubated in M2 medium to resume meiosis. Oocytes developmental capacity was measured by Hoechst 33342 staining after 0 hour, 6 hours and 12 hours of GVBD, which were represented GVBD, MetI and MetII stage of oocytes, respectively. At least 100 oocytes were measured in each group. Unpaired two-tailed Student’s *t*-test. Error bars represent mean ± SD. ***p < 0.001. (D) Wild type and *Ccnb3* mutant oocytes cultured for 4 hours after GVBD used for living cell tracking to observe the spindles formation and first PB extrusion (n = 30 oocytes in each group). Scale bar, 20 μm. (E) Wild type and *Ccnb3* mutant GV-stage oocytes *in vitro* cultured for 6 hours and 12 hours after GVBD were fixed for chromosome spreads (n = 50 oocytes in each group). Scale bar, 5 μm. (F) Immunofluorescence staining for the localization of Rec8 on chromosomes after *Ccnb3* mutant oocytes at GVBD stage were cultured for 6 hours and 12 hours (n = 40 oocytes in each group). Scale bar, 5 μm.

### *Ccnb3* mutation does not affect preimplantation embryonic development and sister chromatid separation

To evaluate the effect of *Ccnb3* mutation on the developmental capacity of oocytes, we injected wild-type sperms into the *Ccnb3*^Δ/Δ^ oocytes through intracytoplasmic sperm injection (ICSI). These restructured *Ccnb3*^Δ/Δ^ embryos could develop into blastocysts with similar efficiency as those embryos in *Ccnb3*^WT/WT^ group (75.6% *vs* 80.6%) (Figure S1F, Table S1), indicating that *Ccnb3* mutation did not affect the cytoplasmic maturation and developmental capacity of embryos. We used ICSI derived blastocysts to establish the embryonic stem cell (ESC) lines (Figure S1G) and analyzed the DNA content and karyotype of the *Ccnb3*^Δ/Δ^ ESCs. We found that the *Ccnb3*^Δ/Δ^ ESCs were triploid (Figure 3A and 3B), which was in agreement with the E7.5 degeneration of the resulted embryos [19] and no separation of homologous chromosome in *Ccnb3*^Δ/Δ^ oocytes.

**Figure 3.**
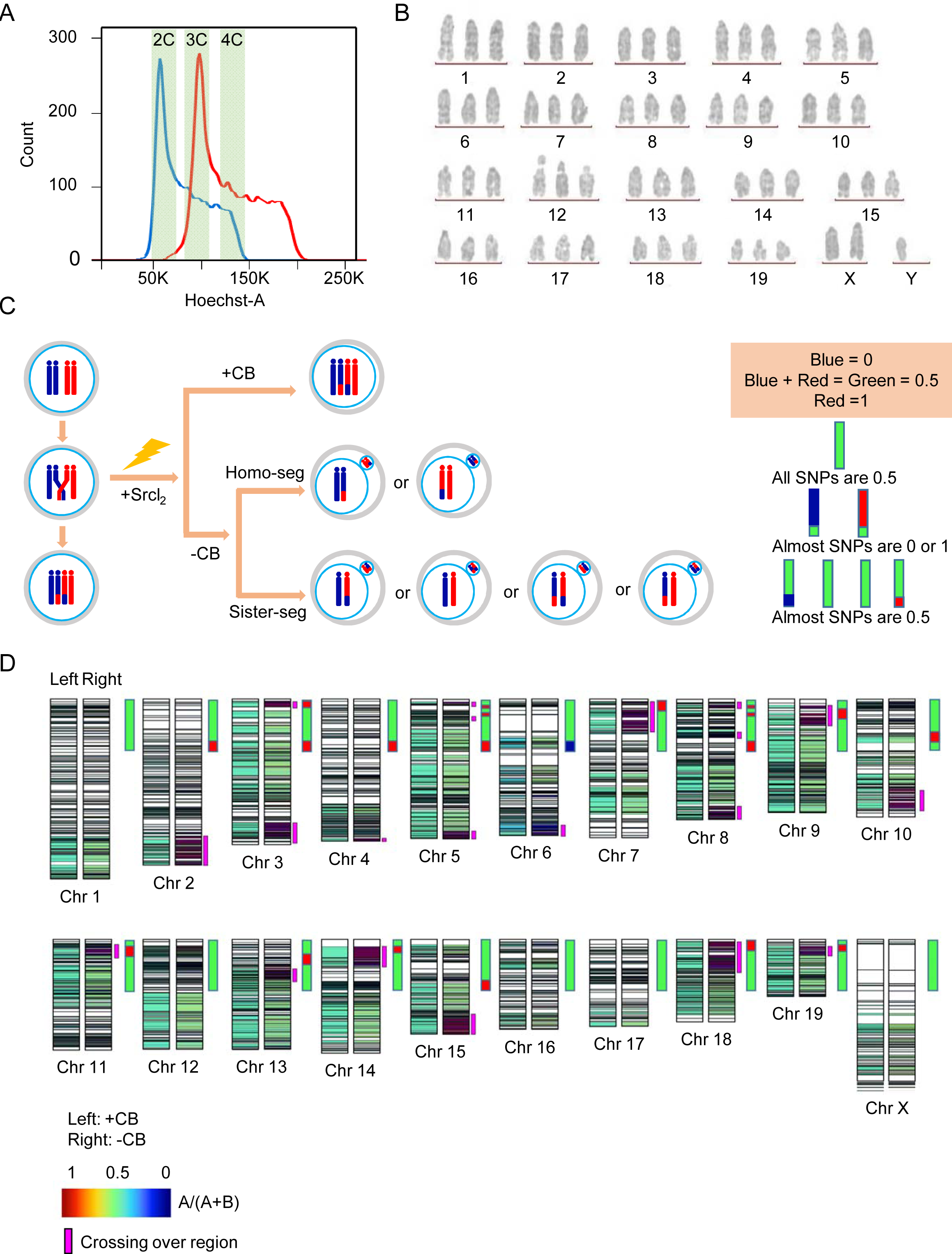
*Ccnb3* mutation did not affect sister chromatid separation after oocyte activation. (A) DNA content analysis of *Ccnb3*^Δ/Δ^ ESCs by FACS. The blue peak represents the wild type ESCs, of which the DNA content were diploid; while the red peak represents the *Ccnb3*^Δ/Δ^ ESCs, of which the DNA content were triploid. 2C, 3C, 4C represents diploid, triploid and tetraploid, respectively. (B) Karyotype analysis of *Ccnb3*^Δ/Δ^ ESCs. *Ccnb3*^Δ/Δ^ ESCs are triploid. (C) Schematic of chromosome segregation during establishment of *Ccnb3*^Δ/Δ^ ESCs by parthenogenetic activation (PA). To distinguish the chromosome segregation happens in homologous chromosomes or the sister chromatids, we established two mESCs from *Ccnb3*^Δ/Δ^ PA embryo, one of which contain 4N chromosomes with no PBs extrusion (with CB treatment, refer to as 4NESCs), the other contain 2N chromosomes with one PB extrusion after PA (without CB treatment, refer to as 2NESCs). Whole genome DNA sequencing was performed. We defined SNPs as 0 (blue) or 1 (red), so if one SNP point is heterozygous, then its value will be 0.5 (blue + red = green), or else its value will be 0 or 1. Thus, the value for all SNPs in 4NESCs would be 0.5. If the segregation happened between homozygous chromosomes, almost SNPs in 2NESCs would be 0 or 1 and the cross-over parts would be 0.5; while if between sister chromatids, almost SNP points would be 0.5, and the cross-over parts would be 0 or 1. (D) Whole genome sequencing analysis of *Ccnb3*^Δ/Δ^ ESCs. The 4NESCs (left) and 2NESCs (right) are sequenced separately. Only the heterozygous sites in 4NESCs and the corresponding sites in 2NESCs are shown. As calculated by the base frequency ratio in one genome locus, the heterozygous sites are marked in range from green to blue and the homozygous sites are marked in red or blue as well as the missing site marked in blank. The homozygous regions are inferred as the crossing-over region through the homozygous sites in 2NESCs.

To distinguish the chromosome segregation between the homologous chromosomes or the sister chromatids, we established two types of ESCs from *Ccnb3*^Δ/Δ^ pathenogenetically activated (PA) embryos, one of which contain 4N chromosomes with no PBs extrusion (with CB treatment, referred to as 4NESCs), the other contain 2N chromosomes with one PB extrusion after PA (without CB treatment, referred to as 2NESCs). Whole genome DNA sequencing was performed. Thus, the value for all Single Nucleotide Polymorphisms (SNPs) in 4NESCs would be heterozygous. If the segregation happened between homozygous chromosomes, SNPs in 2NESCs would be almost homozygous and only the cross-over parts would be heterozygous; while if segregation happened between sister chromatids, SNPs would be almost heterozygous, and the cross-over parts would be homozygous. We confirmed that the sister chromatids of *Ccnb3*^Δ/Δ^ oocytes were separated after PA due to the presence of heterozygous SNPs in whole genome (Figure 3C), which implied that Ccnb3 specifically controlled the meiosis I rather than the meiosis II.

### Ccnb3 is necessary for APC/C activation and MPF activity regulation

To verify the mechanism of regulation defects in *Ccnb3*^Δ/Δ^ oocytes, we examined the activity of MPF and APC/C, which are pivotal in regulating metaphase-anaphase transition. The APC/C controls the metaphase-anaphase progression for which the prerequisites are degradation of Ccnb1 and securin, and inactivation of MPF [20]. To test whether the MetI arrest was caused by the APC/C inactivation in *Ccnb3*^Δ/Δ^ mice, we analyzed the dynamic change of the APC/C substrate securin during *in vitro* maturation. Securin is known for its role in inactivating the cohesin-cleaving enzyme, separase, until the metaphase-anaphase transition [21]. In order to observe the dynamic change of securin, we overexpressed the securin-EGFP by injecting securin-EGFP mRNA into GV-stage oocytes and found that securin-EGFP could not be degraded in the *Ccnb3*^Δ/Δ^ oocytes during meiosis I progression, while the securin-EGFP was degraded in WT oocytes at the time of PB releasing (Figure 4A and 4B). Therefore, *Ccnb3* mutation caused APC inactivation which leads to the MetI arrest. MPF, which is composed of p34^cdc2^ and Ccnb1, promotes entrance into mitotic phase, and its decrease is necessary for anaphase progressing [22, 23]. We detected the MPF concentration by ELISA assay and found that MPF concentration was significantly higher in the *Ccnb3*^Δ/Δ^ oocytes maturation progress, whereas it should be declined at anaphase onset compared with wild type oocytes (Figure 4C). Because the decline of the MPF activity is necessary for metaphase-anaphase transition, we speculated that the high MPF activity hindered the anaphase I onset, and the inactivated APC/C could not decrease the MPF activity which leads to the MetI arrest. To test this hypothesis, we attempted to recover the MetI arrest by modulating the Cdk1 activity. As expected, acute pharmacological inhibition of Cdk1 could partially rescue *Ccnb3* mutation-caused MetI arrest and lead to the extrusion of the first PB (Figure 4D). Our results collectively showed that *Ccnb3* mutation caused APC inactivation and persistence of MPF activity in oocytes, which led to the failure of anaphase I onset.

**Figure 4.**
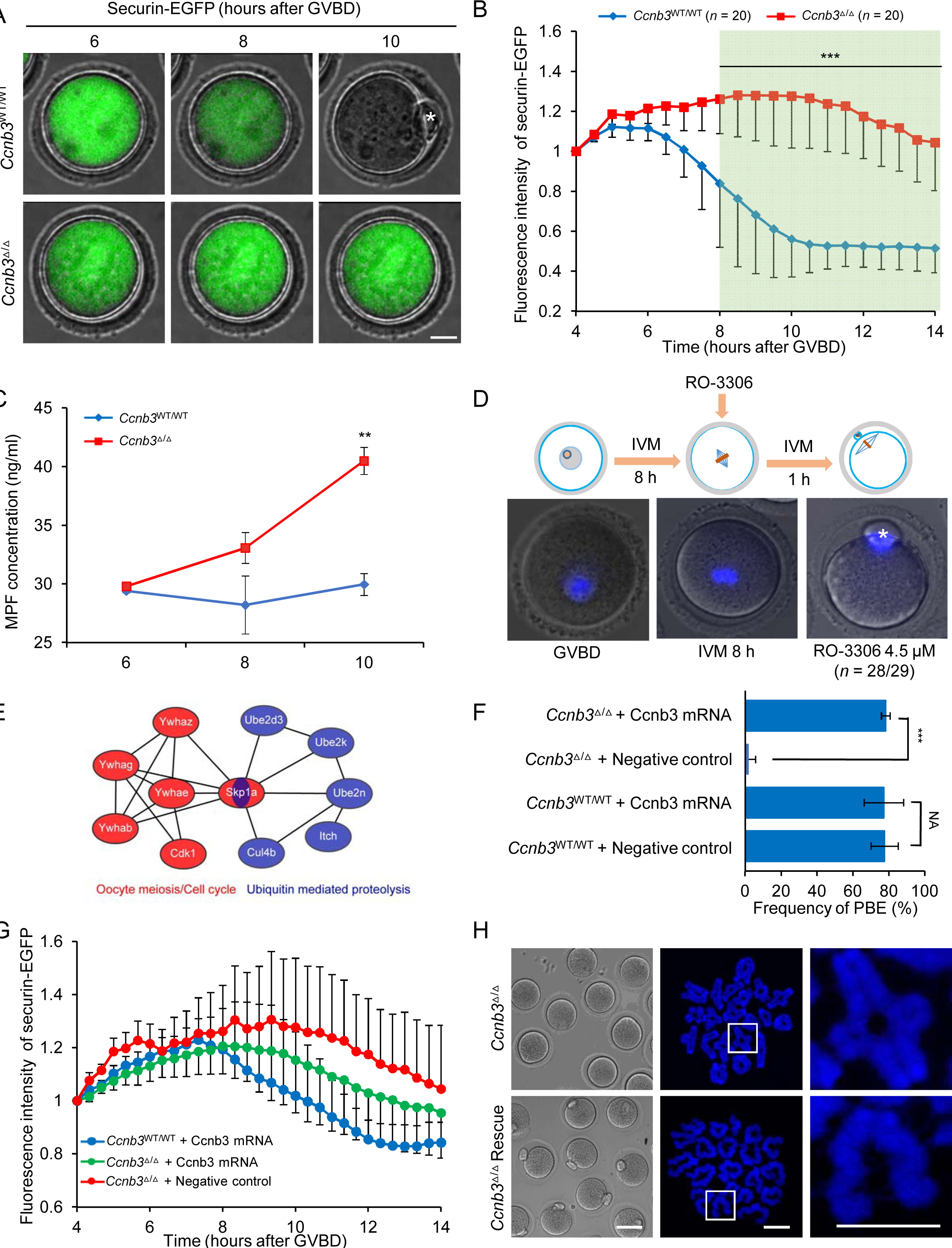
Oocytes with *Ccnb3* mutation fail to initiate anaphase I. (A-B) Time-lapse fluorescence measurement of securin-EGFP expressed from mRNA injected at GV stage. The integrated intensities of securin-EGFP were measured, background corrected, and normalized to the initiative-intensity value obtained per oocyte. GV-stage oocytes were injected with 100 ng/μl securin-EGFP mRNA, incubated in M2 medium containing dbcAMP for 2 hours, then oocytes were washed with dbcAMP-free M2 medium to resume meiosis. At least 20 oocytes were tested in each group. Measurements were aligned to 4 hours after GVBD as the starting time. Scale bar, 20 μm. Statistical analyses for differential securin-GFP fluorescence intensity changes were calculated with two tailed Student’s *t*-test. Error bars represent mean ± standard error of the mean (SEM). ***p < 0.001. (C) Quantification of MPF of oocyte during maturation *in vitro* (n = 40 oocytes). Unpaired two-tailed Student’s *t*-test. Error bars represent mean ± SD. **p < 0.01. (D) Reduction of MPF activity by addition of RO-3306 (a specific inhibitor for Cdk1 activity) in *Ccnb3*^Δ/Δ^ oocyte recovered the MetI arrest. (E) Protein-protein interaction (PPI) relationships of proteins which identified as interacted with Ccnb3 by immunoprecipitation mass spectrometry. These proteins involved in oocyte meiosis or cell cycle and ubiquitin mediated proteolysis pathways. PPI relationships were produced by STRING, and only the connections from databases or validated were shown. (F) Ccnb3 mRNA injection extruded the first PB of MetI arrest oocytes. Unpaired two-tailed Student’s *t*-test. At least 150 oocytes for each group. Scale bar, 40 μm. Error bars represent mean ± SD. ***p < 0.001. (G) Anaphase I initiation was measured by time-lapse fluorescence of securin-EGFP. GV oocytes were injected with 50 ng/μl securin-EGFP mRNA and incubated in M2 medium containing dbcAMP for 2 hours, then oocytes were washed with dbcAMP-free M2 medium to resume meiosis. Scale bar, 40 μm. At least 30 oocytes for each group. NC refers to negative control. (H) Homologous chromosomes were separated after Ccnb3 mRNA injection. *Ccnb3*^Δ/Δ^ oocytes had entered to MetII with univalent sister chromatids after Ccnb3 mRNA injection. At least 80 oocytes were measured. Scale bar, 100 μm.

To search for the proteins that interacted with Ccnb3, we performed the immunoprecipitation mass spectrometry. In total, 174 proteins were identified as interacted with Ccnb3. These proteins enriched in oocyte meiosis or cell cycle (Cdk1, Ywhaz, Ywhag, Ywhab, Ywhae and Skp1a) and ubiquitin mediated proteolysis (Cul4b, Itch, Ube2n, Ube2k, Ube2d3 and Skp1a) pathways (Figure 4E). As known that Cdk1 were reported as the driver of meiosis, the results indicated Ccnb3 directly interacted with Cdk1 and participated into meiosis process.

To investigate whether Ccnb3 protein could recover the MetI arrest in *Ccnb3*^Δ/Δ^ oocytes, we injected Ccnb3 mRNA into oocytes with *Ccnb3* mutation at GV stage (Figure S1H). Our results showed that Ccnb3 mRNA injection could extrude the blocking PB (Figure 4F). By the chromosome spreads assay and the decline of securin-EGFP in Ccnb3 rescue group (Figure 4G and 4H, Figure S1I), we confirmed that MetI arrest caused by *Ccnb3* mutation could be rescued by Ccnb3 mRNA injection, which further proved that Ccnb3 was necessary for APC/C activation.

## Discussion

In this study, we found that *Ccnb3* mutation caused female mouse infertility with the failure of metaphase-anaphase transition in oocyte meiosis I. However, the *Ccnb3* mutant male mice had normal fertility. The reason for female infertility is that Ccnb3 was necessary for APC/C activation to initiate anaphase I. Similar findings were obtained independently with a different targeted mutation in *Ccnb3* [“Cyclin B3 promotes APC/C activation and anaphase I onset in oocyte meiosis” by M.E. Karasu, N. Bouftas, S. Keeney, and K. Wassmann]. The infertility of female mice and normal fertility of male mice with *Ccnb3* mutation suggested that Ccnb3 only functioned in the meiosis in females, which may be related to the long duration of the female meiosis I [1], and reflected different meiosis regulation mechanisms between the two sexes. Interestingly, cyclin A1 is a male mouse specific cyclin whose deletion leads to a block of spermatogenesis before the first meiotic division, whereas female mice were normal [24]. Our results showed that Ccnb3 played an essential role for metaphase-anaphase transition during female meiosis I, consistent with the results in *Drosophila* [25] and *C.elegans* [10]. In addition, the segregation of sister chromatids of *Ccnb3*^Δ/Δ^ oocytes after activation, as well as the cytoplasmic maturation and developmental capacity of *Ccnb3*^Δ/Δ^ embryos, showed no defects, which implied that Ccnb3 is a meiosis-specific required for meiosis I rather than meiosis II.

Successfully execution of metaphase-anaphase transition in eukaryotic cell division requires metaphase cyclins scheduled degradation at different times, especially cyclin B. There are at least three types of cyclin B (B1, B2 and B3) in mammals and it appears that Ccnb1 is primarily responsible for MPF activity [26]. Only two B-type cyclins, Ccnb1 and cyclin B2 (Ccnb2), have been intensively studied so far in mammals. Mouse lacking Ccnb1 could not viable, whereas *Ccnb2*-null mouse had no apparent defect [27]. Our results showed that Ccnb3 played an essential role for metaphase-anaphase transition during female meiosis I and enriched the knowledge of cyclin B family.

The metaphase-anaphase transition during meiosis I takes place when the activity of MPF reduced and the APC/C activated. MPF activity is critically regulated during the meiosis and APC/C trigger the degradation of key cell cycle regulators [20, 28]. Securin and Ccnb1 are the two established substrates for APC/C whose degradation releases separase and inactivates Cdk1 at the metaphase-anaphase transition [20]. We found that the *Ccnb3* deficient oocytes maintained high level of MPF activity which was probably due to high Cdk1 activity. Adding an inhibitor of Cdk1 restored the metaphase arrest and released the PBs, suggesting that Cdk1 activity needs to be efficiently reduced in oocytes for homologous chromosome segregation. Securin was not degraded during *in vitro* maturation suggested that APC/C-mediated MPF reduction is necessary for metaphase-anaphase transition. We also found the interaction between Ccnb3 and Cdk1. These observations implied that Ccnb3 may form a MPF complex with Cdk1 and participate the regulation of MPF and APC/C activity [29].

Although we have elucidated the role of Ccnb3 in female meiosis, the direct target of Ccnb3 have not been found yet. In addition, although core regulatory machinery of metaphase to anaphase transition in female meiotic cells and mitotic cells is similar, Ccnb3 may play different roles in them. Our study which reveals that Ccnb3 serve as a female meiosis-specific cyclin for metaphase-anaphase transition in meiosis I opens an avenue to elucidate the cell cycle regulation mechanisms for meiosis in mammals.

## STAR★Methods

### Contact for Reagent and Resource Sharing

Further information and requests for resources and reagents should be directed to and will be fulfilled by the Lead Contact, Wei Li.

### Experimental Model and Subject Details

#### Experimental Animals

Specific-pathogen-free (SPF)-grade mice were obtained from Beijing Vital River Laboratories and housed in the animal facilities of the Chinese Academy of Sciences. All the studies were carried out in accordance with the Guidelines for the Use of Animals in Research issued by the Institute of Zoology, Chinese Academy of Sciences. The mice used in this study were CD-1 strains and the genome editing was carried out by CRISPR/Cas9. Six sgRNAs were designed for targeting different sites of the mouse *Ccnb3* gene (sgRNA-*Ccnb3*-1∼6). The efficiency of sgRNAs was verified by mouse fibroblast cell transfection, cell sorting, subsequent PCR identification, T7 endonuclease I (T7EI) assay and Sanger sequencing. SgRNA-*Ccnb3*-1 showed the highest mutation ratio (12/20, 60%) according to T7 EI and Sanger sequencing, which was selected for *in vitro* transcription and injection. The injection concentration by intracytoplasmic sperm injection (ICSI) of Cas9 mRNA/sgRNA-*Ccnb3*-1 was 100:50 ng/μl. The biallelic mutations of *Ccnb3* in mouse embryos were produced with relatively high efficiency via zygote injection. Finally, we screened out the Δ29 bp mice for the next experiment. All sgRNAs were listed in the Supplementary information, Table S2.

## Method Details

### RNA extraction and qRT-PCR

Total RNA was extracted with TRIzol reagent (Invitrogen, 15596-018) from 20 oocytes in each group. Real-time PCR was carried out with THUNDERBIRD SYBR qPCR Mix (TOYOBO) using total cDNA as the template, primer, and ROX (TOYOBO) in a total volume of 20 μl. The thermal cycling conditions of conventional real-time PCR were 1 min at 95 °C, followed by 45 cycles of 15 s at 95 °C, 15 s at 60 °C, and 45 s at 72°C, and the melting curve were 1 min at 95°C, 30 s at 60°C, 30 s at 95°C. All reactions were performed using an Agilent Technologies Stratagene Mx3005P Real-time PCR System (Applied Biosystems). Relative gene expression was analyzed based on the 2-ΔΔCt method with Actin as internal control. At least three independent experiments were analyzed. All primers were listed in the Supplementary information, Table S3.

### Histological analysis

Ovaries from wild type and *Ccnb3* mutant female mice were fixed in 4% paraformaldehyde overnight at room temperature. The ovaries were dehydrated stepwise through an ethanol series (70%, 80%, 90%, 100% ethanol) and processed for paraffin embedding. 5 µm sections were cut with a Leica slicing machine (Leica RM2235, Leica Biosystems, Germany) and mounted on poly-D-lysine coated glass slices (Zhong Shan Golding Bridge biotechnology, Beijing, China). After dewaxing and hydration, the sections were stained with H&E using standard methods and imaged with a Leica Aperio VERSA 8 microscope (Leica Biosystems, Germany).

### *In Vitro* maturation of GV-stage oocytes

We isolated GV stage oocytes from minced ovaries of 8- to 10-week-old CD-1 female mice. Oocytes were cultured in M2 medium for at least 12 hours. The culture was conducted in an incubator under environmental conditions of 5% CO_2_, 37°C, and saturated humidity. To evaluate the developmental capacity of oocytes, oocytes were stained with Hoechst 33342 at different time during *in vitro* culture as figure legends showed. For the rescue experiment, Oocytes were cultured in medium containing RO3306 at different concentrations and for different lengths of time as indicated in the figure legends.

### Time-lapse confocal microscopy of live oocytes

The fluorogenic, cell-permeable reagent SiR-Tubulin (Cytoskeleton, CY-SC002) was used to image tubulin in living cells. GV-stage oocytes were injected with 10 pl of 100 ng/μl mRNA encoding securin-EGFP in M2 medium containing 250 mM dbcAMP using methods described elsewhere [30]. Following mRNA injection, oocytes were cultured for 2-3 hours at 37°C to allow securin-EGFP expression. Oocytes were cultured in dbcAMP-free M2 medium placed in an EMBL environmental microscope incubator (EMBL, GP106), allowing cells to be maintained in a 5% CO_2_ atmosphere at 37°C with humidity control during imaging. Time-lapse image acquisitions were performed using a customized Zeiss LSM510 META confocal microscope equipped with a 532 nm excitation laser, a long-pass 545 nm emission filter, a 403 C-Apochromat 1.2 NA water immersion objective lens (Carl Zeiss), and an in-house-developed 3D tracking macro [31].

### Preparation and staining of chromosome spreads

Chromosome spreads of mouse oocytes during meiotic maturation were prepared using methods previously described [32]. Briefly, oocytes were exposed to acid Tyrode’s solution (Sigma) to remove the zona pellucida under the microscope to avoid over-digestion. After a brief recovery in M2 medium, the oocytes were transferred onto glass slides and fixed in a solution of 1% paraformaldehyde in distilled H_2_O (pH 9.2) containing 0.15% Triton X-100 and 3 mM dithiothreitol. The dried chromosome spreads were stained with 10 μg/ml Hoechst 33342 for DNA counterstaining, following immunofluorescence staining when required. The slides were dried slowly in a humidified chamber for several hours, and then blocked with 2% BSA in PBS for 1 hour at room temperature or overnight at 4°C and incubated with primary antibodies overnight at 4°C. After brief washes with washing buffer, the slides were then incubated with corresponding secondary antibodies for 1 hour at room temperature. Rabbit anti-Rec8 antibody (Abcam, ab192241) for marking homologous chromosomes was used as primary antibodies, and appropriate secondary antibodies conjugated with Alexa Fluor Cy3 (Invitrogen) was used. The samples were observed under a laser-scanning confocal microscope (Zeiss LSM 780, Germany).

### *Ccnb3*^Δ**/**Δ^ ESCs derivation and cell culture

CD-1 mouse blastocysts were used to derive ESCs. The culture medium composed of N2B27 medium (Gibco), 1 mM L-glutamine, 0.1 mM β-mercaptoethanol, 50 U/ml penicillin, 50 μg/ml streptomycin, 3 μM CHIR99021 (Stemgent), 1 μM PD0325901 (Stemgent) and mLIF (ESGRO). The embryos were incubated at 37°C in a 5% CO_2_ incubator for 4-5 days, and then the formed outgrowths (named passage 0) was picked separately by a glass pipette and dissociated with digest solution (0.25% trypsin-100 μM EDTA). The cells replanted and survived from the first trypsinization of the outgrowth were counted as passage 1 (P1). For routine passage, the passage ratio was about 1:8, and mouse ESCs were passaged around every 3 days.

### DNA content analysis of *Ccnb3*^Δ**/**Δ^ ESCs

*Ccnb3*^Δ/Δ^ ESCs were purified and analyzed by FACS. Briefly, single-cell suspensions were obtained by trypsin-EDTA digestion and repetitive pipetting and sieved through a 40-mm cell strainer. Cells (1C) were incubated with 10 μg/ml Hoechst 33342 (Invitrogen, H3570) for 15-20 min at 37°C before analysis. Data were collected with a BD FACS Aria II cell sorter (BD Biosciences) and MoFlo XDP cell sorter (Beckman-Coulter). *Ccnb3*^WT/WT^ ESCs (2C) were used as a control. The percentage of haploid cells was calculated based on the percentages of 1C peak (including haploid cells at G1 phase), 2C peak (including haploid cells at G2/M phase and diploid cells at G1 phase), and 4C peak (including diploid cells at G2/M phase).

### Karyotype analysis of *Ccnb3*^Δ/Δ^ ESCs

*Ccnb3*^Δ/Δ^ ESCs were incubated with 0.2 mg/ml nocodazole (Sigma, M1404) for 3 hours. After trypsinization, the ESCs were suspended in 0.075 M KCl at 37°C for 30 min. Then, the cells were fixed with solution consisting of methanol and acetic acid (3:1 in volume) for 30 min and then were dropped onto pre-cleaned slides. The cells were stained with Giemsa stain (Sigma, GS500ML) for 15 min after being incubated in 5 M HCl. More than 30 metaphase spreads were analyzed.

### DNA sequencing of *Ccnb3* PA derived ESCs

Collected oocytes were parthenogenetic activated in Ca^2^+-free CZB containing 10 mM Sr^2^+ with (5 nmol) and without Cytochalasin B (CB) for 6 hours. Parthenogenetic activated embryos were used to establish ESCs. Two mESCs from *Ccnb3*^Δ/Δ^ PA embryo were established, one of which contain 4N chromosomes with no PBs extrusion (+CB, 4NESCs), the other contain 2N chromosomes with one PB extrusion after PA (-CB, 2NESCs). At least 1 × 10^7^ ESCs were collected for DNA sequencing. DNA was isolated and checked by using the Nano Photometer Spectrophotometer and Qubit 2.0 Flurometer. Used 1 μg gDNA template according to TruSeq DNA Sample Preparation Guide (Illumina, 15026486 Rev.C) method and process for library preparation. Used Illumina HiSeq X Ten, PE150 strategy to sequencing the identified library. Used Illumina HiSeq X Ten, PE150 strategy to sequencing the identified library. The clean reads were mapped to the mouse reference genome (mm9) using a Burrows–Wheeler Aligner (BWA, version 0.7.15) [33]. After removing duplicated reads, the base distribution for each chromosomal location was calculated using the pysamstats (version 1.0.0) (https://github.com/alimanfoo/pysamstats). The single nucleotide variations (SNVs) in 4NESCs sample with more than 10X coverage were used for the SNP identification. In briefly, the sites were picked for analysis with covered by two type bases and located within non-repeat genome regions (http://repeatmasker.org/). Then, the sites with minor base taking more than 30% reads were used for the analysis. Next, we calculated the corresponding base frequency distribution in the 2NESCs sample. If the homologous chromosome separation happening, the 2NESCs sample showed the homozygous state in the identified SNV sites; whereas, the cell showed sister chromatid separation and the homologous chromosomes were retained.

### MPF kinase assay

Forty oocytes incubated in 200 μl PBS were used in each group and lysised by repeated freezing and thawing (Freeze 30 min in liquid nitrogen, thaw at 37°C, repeated 5 times). MPF kinase assay was carried by using Mouse MPF ELISA Kit (JiangLai Biology) according to the manufacturer’s instruction. Each sample makes 3 repetitions. Except for blank hole, 100 μl enzyme reagent was added to each hole. Then the plate was placed at 37°C for 60 min. Discarded the liquid and washed the plate 5 times with 1 × washing liquid. Added 50 ul chromogenic agent A then add 50 μl chromogenic agent B to each hole, gently oscillated and mixed together, avoid light at 37°C for 15 min. Added 50 μl terminating solution to each hole to terminate the reaction. Zero with a blank hole, measured the absorbance (OD) of each hole at 450 nm wavelength within 15 min. Used the concentration of standard products as a horizontal coordinate and OD of sample as a vertical coordinate to make a standard curve.

### Immunofluorescence analysis

Oocytes were first fixed in 4% paraformaldehyde at room temperature for 30 min, and then permeabilized in PBS containing 0.5% Triton X-100 for 20 min. Next, oocytes were blocked in PBS with 2% BSA for 1 hour, and then incubated with FITC-α-tubulin (F2618, Sigma) first antibody at room temperature for 1 hour. After washing 3 times with PBS, DNA was stained with Hoechst 33342. The samples were then mounted on slides and were observed under a laser-scanning confocal microscope (Zeiss LSM 780, Germany). At least 40 oocytes were examined in each treatment, and each treatment was repeated three times.

### Immunoprecipitation and mass spectrometry

Magnetic Beads Protein G were coated with 5 μg of primary antibody in IP wash buffer (50 mM Tris-HCl, pH 7.4, 150 mM sodium chloride, 1 mM MgCl_2_, and 0.05% NP-40) containing 5% BSA overnight with rotation at 4°C. Then, we collected approximately 2000 MII oocytes, and added 100 μl of IP lysis buffer (150 mM NaCl, 50 mM Tris-HCl pH 7.4, 1 mM EDTA, 0.1% SDS, 1% NP-40, 0.5% sodium deoxycholate, 0.5 mM DTT, 1 mM PMSF/cocktail) followed by incubation on ice for 10 min. Next, we centrifuged the IP lysate at 14,000 rpm for 10 minutes at 4°C, removed 100 μl of the supernatant and added this to 900 μl of beads-antibody complex in IP Immunoprecipitation Buffer (860 μl IP wash buffer, 35 μl 0.5 M EDTA), and incubated this with rotation overnight at 4°C. After washing, the immunoprecipitate was added 50 μl elution buffer and the supernatant was used for mass spectrometry analysis. Mass spectrometry analysis was performed by BGI Company.

### Quantification and Statistical Analysis

Statistical parameters including statistical analysis, statistical significance and n value are reported in the Figure legends. Statistical analyses were performed using Prism Software (GraphPad). For statistical comparison, Student’s *t*-test was employed. A value of p < 0.05 was considered significant.

## Acknowledgments

We thank Lijuan Wang for her help of live cell imaging and Shiwen Li, Xili Zhu for the help with immunofluorescent staining. This study was supported by the National Key Research and Development Program (2017YFA0103803), the China National Postdoctoral Program for Innovative Talents (BX201700243 to L.W.), the National Natural Science Foundation of China (31621004 and 81571356), and Key Research Projects of the Frontier Science of the Chinese Academy of Sciences (QYZDY-SSW-SMC002 to Q.Z.).

## Author Contributions

W.L., Q.Z., and Y.Z. conceived of and designed the experiments; L.W. performed the ICSI, L.Z. performed the qPCR, genotyping and MPF kinase assay; J.W. performed the protein mass spectrometry; G.F. analyzed the protein spectrum data and DNA sequencing data; other experiments and analysis were performed by Y.L. and Z.H.; Y.L wrote the manuscript with the help of other authors.

## Supplemental Information Titles and Legends

Figure S1

Supplemental Figure legends: Figure S1. Characters of *Ccnb3* mutant mice

Table S1. Developmental capacity of *Ccnb3*^Δ/Δ^ oocytes after ICSI

Table S2. Related to experimental procedures–sgRNAs designed in the study

Table S3. Related to experimental procedures–primers used in the study

